# Using rapid invisible frequency tagging (RIFT) to probe the attentional distribution between speech planning and comprehension

**DOI:** 10.1101/2024.04.17.589897

**Authors:** Cecília Hustá, Antje Meyer, Linda Drijvers

**Author notes:** C. Husta Wundtlaan 1 6525 XD NIJMEGEN NETHERLANDS.

## Abstract

Interlocutors often use the semantics of comprehended speech to inform the semantics of planned speech. Do representations of the comprehension and planning stimuli interact on a neural level? We used rapid invisible frequency tagging (RIFT) and EEG to probe the attentional distribution between spoken distractor words and target pictures in the picture-word interference (PWI) paradigm. We presented participants with auditory distractor nouns (auditory (f1); tagged at 54Hz) together with categorically related or unrelated pictures (visual (f2); tagged at 68Hz), which had to be named after a delay. RIFT elicits steady-state evoked potentials, which reflect attentional allocation to the tagged stimuli. When representations of the tagged stimuli interact, integrative effects have been observed at the intermodulation frequency resulting from an interaction of the base frequencies (f2±f1; Drijvers et al., 2021). Our results showed clear power increases at 54Hz and 68Hz during the tagging window, but no differences between related or unrelated conditions. More interestingly, we observed a larger power difference in the unrelated compared to the related condition at the intermodulation frequency (68Hz – 54Hz: 14Hz), indicating stronger interaction between the auditory and visual representations when they were unrelated. Our results go beyond standard PWI results (e.g., Bürki et al., 2020) by showing that participants do not have more difficulty visually attending to the related pictures or inhibiting the related auditory distractors. Instead, processing difficulties arise when the representations of the stimuli interact, meaning that participants might be trying to prevent integration between the auditory and visual representations in the related condition.

**Significance statement:** Studying speech planning during comprehension with EEG has been difficult due to a lack of appropriate methodology. This study demonstrates that rapid invisible frequency tagging (RIFT) can explore attentional allocation to speech planning and comprehension stimuli, as well as their interaction. Our results show that the content of the speech planning and comprehension representations affects their interaction in the neural signal, which should always be considered when these processes are studied jointly. In future work, RIFT could be used to investigate speech planning and comprehension in more conversational settings, as tagging can be added to videos or speech segments. This is the first study that demonstrates that RIFT can be used together with EEG to study cognitive phenomena.

## 1. Introduction

A robust finding of picture-word interference (PWI) studies is the semantic interference effect: slower responses when naming target pictures that are presented with categorically related than with unrelated auditory or written distractors (for review see Bürki et al., 2020). The PWI literature usually focuses on the locus of the interference effect, arising from competition between related words during lexical selection (e.g., Schriefers et al., 1990) or from the need to remove related distractors from the response buffer (e.g., Finkbeiner & Caramazza, 2006). Given that the representations of targets and distractors compete on some level, they must also interact. In this study, we leverage the PWI paradigm to better understand comprehension, speech planning, and their interaction, as in this paradigm participants simultaneously comprehend auditory distractors while preparing to name picture targets. We will investigate how participants distribute their attention to representations of speech planning and comprehension stimuli. Crucially, we will also look at how these representations interact in the neural signal.

Speech planning as well as listening requires attentional capacity, which means that attentional resources are shared between speech planning and comprehension when they take place concurrently. In line with this, evidence from dual-tasking studies shows that there is mutual interference between speech planning and comprehension (Bögels et al., 2015; Daliri & Max, 2016; Fargier & Laganaro, 2016, 2019). Does the content of the speech planning and comprehension affect the attentional distribution? In many conversations, the planned utterance depends on the content of comprehension. This means that interlocutors use the semantics of comprehended speech to inform the semantics of speech planning (Bögels et al., 2015, 2018). Thus, relatedness of speech planning and comprehension streams could affect how future speakers distribute their attentional resources.

There is some evidence that the comprehension and production system interact. Several theories assume that the two systems work jointly to improve their functioning (Chang, 2002; Federmeier, 2007; Pickering & Gambi, 2018; Rohde, 2002). For example, the production system can aid with predicting upcoming words during comprehension. The overlap and potential interaction of representations within the comprehension and production system is understood less, as it is difficult to study the interaction between covert and complex processes.

A novel approach, rapid invisible frequency tagging (RIFT), has recently been put forward for investigating both attentional allocation to multiple stimuli and stimulus interaction (Brickwedde et al., 2022; Duecker et al., 2021; Ferrante et al., 2023; Pan et al., 2021; Seijdel et al., 2023, 2024; Zhigalov et al., 2019, 2021; Zhigalov & Jensen, 2020). In this method the luminance of the visual stimuli and the amplitude of the auditory stimuli is periodically modulated at high frequencies (>50Hz), which leads to imperceptible tagging that does not interfere with endogenous oscillations (Minarik et al., 2023). Visual and auditory tagging produce robust steady-state evoked potentials (SSEP). The strength of these potentials reflects visual or auditory attention towards the tagged stimuli (Toffanin et al., 2009). Tagging at two different frequencies also creates increased power at intermodulation frequencies, which results from the nonlinear interaction of the base frequencies ((f2±f1); e.g., Regan et al., 1995). The power at the intermodulation frequency is thought to reflect the strength of interaction between the representations of the two stimuli (Drijvers et al., 2021; Seijdel et al., 2024). In this study, we utilized the RIFT approach to study interaction between speech planning and comprehension.

To study speech planning during comprehension, we utilized a delayed-naming PWI paradigm combined with RIFT and EEG. In this study, participants named visually tagged pictures (68Hz luminance modulation) that were presented simultaneously with auditorily tagged distractor words (54Hz amplitude modulation). We examined the SSEPs at the tagging frequencies to determine whether relatedness affects how participants distribute their attentional resources to speech planning stimuli (i.e., target picture) and to comprehension stimuli (i.e., distractors). Importantly, we also examined the intermodulation frequency to determine whether the participants’ representations of the speech planning and comprehension stimuli interacted differently depending on their relatedness.

## 2. Materials and methods

The present study was approved by the Ethics Committee of the Social Sciences department of the Radboud University Nijmegen (ECSW-2019-019).

## 2.1. Participants

Thirty-one, right-handed, native Dutch-speaking participants with no hearing or language impairments and normal or corrected to normal vision took part in the experiment for financial compensation. Data of one participant were excluded because there were too many physiological artifacts. The remaining 30 participants had a mean age of 23.13 years (range: 18 - 32) and 11 were male.

### 2.2. Materials

For the pretest, 52 categorically related picture distractor pairings were inspired by previous studies (Mahon et al., 2007; Schriefers et al., 1990) as well as newly created (see Supplementary Materials). The present study needed longer distractors than standard PWI studies to ensure the speech input was long enough (>700ms) for auditory tagging. All distractors were pre-recorded by a female speaker using a slower pace using 3.0.0 Audacity(R). The log 10-word frequencies (WF) for the distractors and picture names were calculated using SUBTLEX-NL (Keuleers et al., 2010). The mean WF of the distractors was 2.09 (range: 0.48 – 3.59); for the picture names it was 2.79 (range: 1.15 – 5.11). The distractors were paired with different pictures from our set to create the unrelated picture distractor pairs. This was done to find pairings with related and unrelated distractors of similar length and WF per picture. Thus, each picture was paired with one categorically related and one unrelated distractor. The pictures were taken from the BOSS photograph database (Brodeur et al., 2014) as well as online.

The PWI materials were tested in a behavioral pretest with 14 different native Dutch participants. In the pretest participants were familiarized with the target picture names and the auditory distractors. They read through all the picture names, which were printed underneath the pictures and they listened to a recording of all of the distractors in a random order. In the pilot experiment, each target-distractor pair was presented twice. Participants first saw a fixation cross for 700ms, after which the target picture appeared simultaneously with the auditory distractor. Participants’ task was to name each picture as fast as possible.

Trained research assistants transcribed the responses and determined the naming latencies using Praat (Boersma, P. & Weenink, D., 2023). They were blind to the experimental conditions. We excluded incorrect responses and responses with naming latencies 2.5 SD from the mean per condition. Naming latencies were analyzed with linear mixed-effects model fitted with lmerTest package (version 3.4; including the lme4 package, Bates et al., 2015) with random intercepts for participant, item, and picture category. As expected, the naming latencies were shorter in the unrelated condition (*M* = 721.47, *SD* = 165.44) than in the related condition (*M* = 738.70, *SD* = 181.94, β = 18.17, S.E. = 4.95, t = 3.67, p < 0.001).

Based on the pretest, we selected 32 targets that showed the strongest interference for the EEG experiment. The average naming latencies for this subset was 718.76ms (*SD* = 161.47) in the unrelated condition and 739.74ms (*SD* = 181.96) in the related condition. The mean duration of the distractors was 876.88ms (range 722 – 1044). Each target-distractor pair was presented twice during the EEG experiment, with at least ten intervening trials. We created 30 pseudorandomized lists with MIX (Van Casteren & Davis, 2006). The same condition occurred maximally on four consecutive trials.

### 2.3. EEG acquisition

EEG was recorded using 32-scalp electrodes using actiCap system (Brain Products, Germany) arranged according to the 10-20 system. Thirty-one electrodes were positioned on the scalp and one electrode was placed on the left mastoid. The right mastoid was used as an online reference. The data was recorded using a 1000Hz sampling rate. We adjusted the impedances of all electrodes below 10 kΩ.

### 2.4. Procedure

The participants were first familiarized with the picture names, by looking at each picture and reading its associated name on a printed sheet. All of the experimental stimuli were presented using MATLAB 2023b (Mathworks Inc, Natrick, USA) and the 3.0.17 Psychophysics Toolbox (Brainard, 1997; Mario Kleiner et al., 2007). The stimuli were projected with a PROPixx DLP LED projector (VPixx Technologies Inc., Saint-Bruno-de-Montarville, Canada), and using a GeForce GTX960 2GB graphics card with a refresh rate of 120Hz. This setup could achieve a presentation rate up to 1,440Hz, as the projector interprets the four quadrants and three-color channels of the GPU screen buffer as frames, that can be projected in rapid succession (4 quadrants * 3 color channels * 120HzL=L1,440Hz).

The experiment took place in a dimly-lit room. During the task, participants first saw a fixation cross presented together with a beep for 200ms, which was followed by the baseline period of 1000ms, during which participants only saw a fixation cross. Then participants saw a picture for 1000ms (i.e., tagging window). The luminance of the pixels of this picture were multiplied with a 68Hz sinusoid (i.e., visual frequency tagging). Participants were instructed to prepare the picture names as soon as possible in order to speed up naming after the response signal. The auditory distractors were presented simultaneously with the pictures. The amplitude of the auditory distractors was multiplied with a 54Hz sinusoid (i.e., auditory frequency tagging). Subsequently, participants saw a gray square for 400ms, after which an exclamation mark appeared for 1600ms. Participants were instructed to name the picture as fast as possible after the onset of the exclamation mark. After that, participants could blink at the onset of the three stars (see Figure 1).

**Figure 1.**
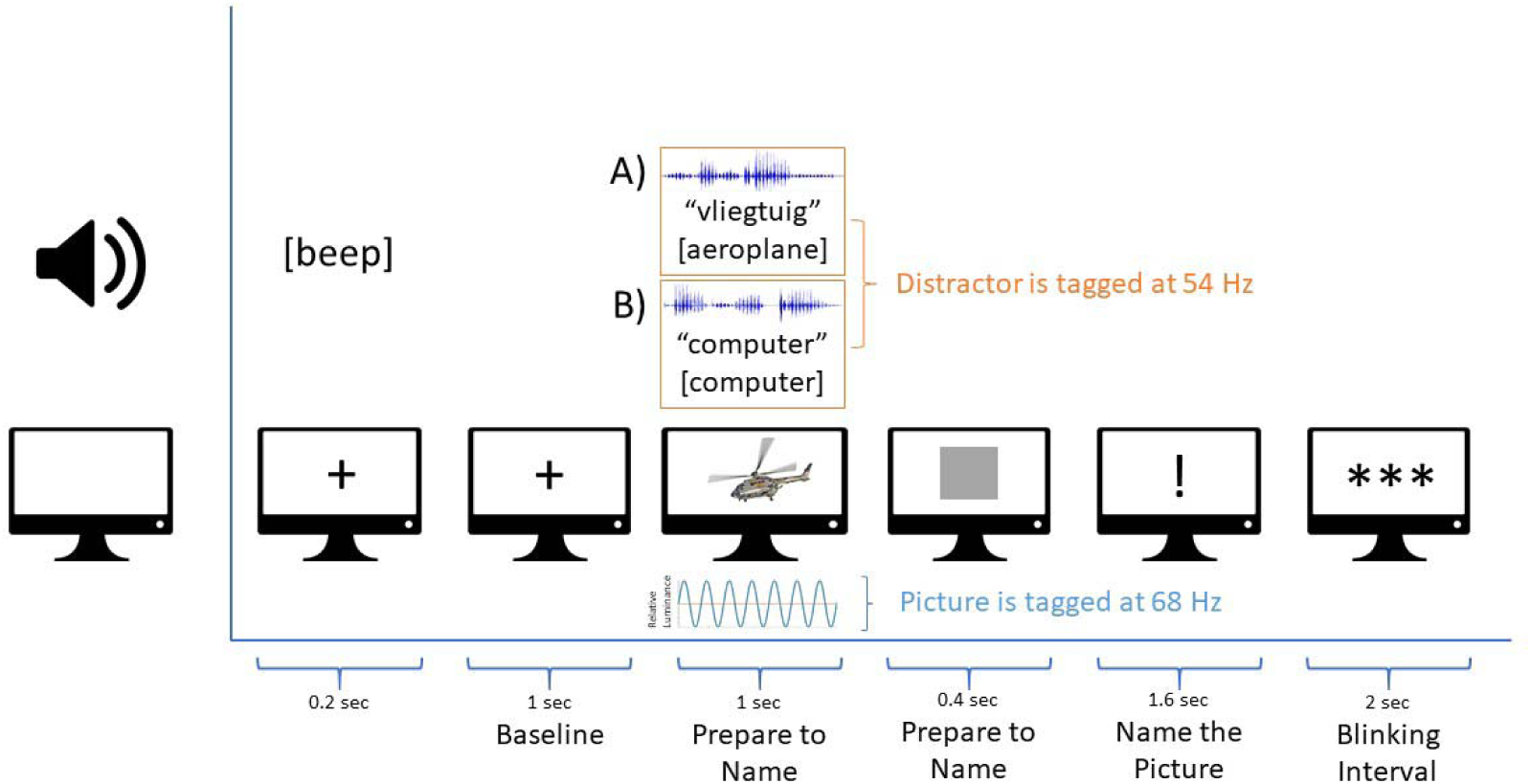
Experimental Paradigm. Participants saw pictures with luminance modulations of 68Hz while listening to auditory distractors that included amplitude modulations at 54Hz. The auditory distractors were either A) categorically related or B) unrelated to the target pictures. Participants were asked to prepare the picture names as soon as possible, but only to say them at loud at the onset of the exclamation mark.

### 2.5. EEG preprocessing

We analyzed all EEG data using FieldTrip (Oostenveld et al., 2010) in Matlab (R2020b). The data was segmented from 1000ms before to 1000ms after target picture onset. Channels with excessive noise were removed (no more than four channels were excluded per participant). All channels were re-referenced to an average mastoid reference. We filtered the data with a band-pass filter between 0.1 and 100Hz, and a notch filter at 50Hz, 100Hz, and 150Hz to remove the line noise and its harmonics. We visually inspected the data and removed the segments containing non-physiological artifacts, resulting from other devices or electrical phenomena. Independent component analysis (ICA) was performed to correct for EOG artifacts and steady muscle activity. The remaining physiological artefacts were removed manually. The individual removed EEG channels were interpolated by a weighted average of the data from neighboring channels of the same participant. An average of 55.22 trials (min: 48, max: 63) were included per participant and per condition.

### 2.6. Analysis

The behavioral responses were scored as correct if participants named the picture with the familiarized name or its synonym. Omissions, names of other targets, or early responses were excluded from all analyses (1.07% of the trials). The naming latencies from the EEG experiment were determined based on an automatic threshold, as we did not expect any differences between conditions in a delayed naming experiment. We excluded responses with naming latencies 2.5 SD from the mean per condition for the RT analysis. The naming responses were analyzed using a linear mixed-effects model fitted with lmerTest package (version 3.1-3; including the lme4 package, Bates et al., 2015) with random intercepts for participants.

To examine success of the frequency tagging at the base frequencies we performed both power and coherence analysis. The coherence analysis did not show different results to the power analysis; thus, we only describe it in the Supplementary Materials. The power spectra were calculated per participant and per condition between 1 and 80Hz using 1Hz steps for the tagging interval as well as the baseline. Each frequency step was modulated with a boxcar taper, followed by the Fourier transform of the tapered signal. We computed averages per condition for every participant.

We first evaluated power differences between the tagging window and the baseline at the main tagging frequencies (i.e., 68Hz and54 Hz) and at the intermodulation frequency (i.e., 14Hz) using two-tailed non-parametric cluster-based permutation tests (Maris & Oostenveld, 2007). This test computed dependent samples t-tests between the conditions for the frequency of interest and for all electrodes. If two or more neighboring electrodes reached significance, they formed a cluster. Subsequently, a sum of all the t-values of each cluster was calculated. To control for the family-wise error rate, the two conditions were combined and subsequently randomly separated into two artificial groups 5000 times. The summed t-values of all of the randomly generated clusters were computed in the same way as described above. Subsequently, a distribution of these summed t-values was generated. Comparing the summed t-values computed based on the original clusters to this distribution resulted in Monte-Carlo significance probabilities, which were considered significant if smaller than 0.05. The differences between related and unrelated condition were examined only after the tagging peaks significantly differed from the baseline.

To compare power differences between the conditions, we first computed percentage power change between the tagging window and the baseline. The percentage power change was calculated as ((Pow during tagging window. – Pow during baseline) / Pow during baseline) *100. The condition differences were also tested with non-parametric cluster-based permutation tests using the same procedure described above.

## 3. Results

### 3.1. Behavioral analysis

Less than 2% of responses were marked as incorrect and excluded from all further analyses. In line with previous studies that used the PWI paradigm with delayed naming (Mädebach et al., 2011; Piai et al., 2011), we found that the naming latencies did not significantly differ in the related (*M* = 488.52, *SD* = 188.02) as compared to the unrelated condition (*M* = 498.33, *SD* = 184.45, β = -8.31, S.E. = 4.90, t = -1.70, p = 0.090). This indicates that participants started speech planning before the response period.

### 3.2. Main frequency-tagging peaks

We first examined whether the visual and auditory SSEP were larger in the tagging window compared to the baseline. The nonparametric cluster-based permutation test (for both conditions combined) showed that the visual SSEP arising from the tagging at 68Hz resulted in a significant power peak in the tagging window compared to the baseline (p < 0.001). Based on visual inspection about 63% of the participants showed the visual tagging peak, similar to what has been reported for MEG data in Pan et al., (2021). The auditory SSEP arising from the tagging at 54Hz also showed significant power peak in the tagging window compared to the baseline (p < 0.001). Based on visual inspection about 77% of the participants showed the auditory tagging peak. The visual SSEPs were significantly different from baseline mainly in the occipital and central electrodes, while the auditory SSEPs were significantly different from baseline in most electrodes (see Figure 2A.).

**Figure 2.**
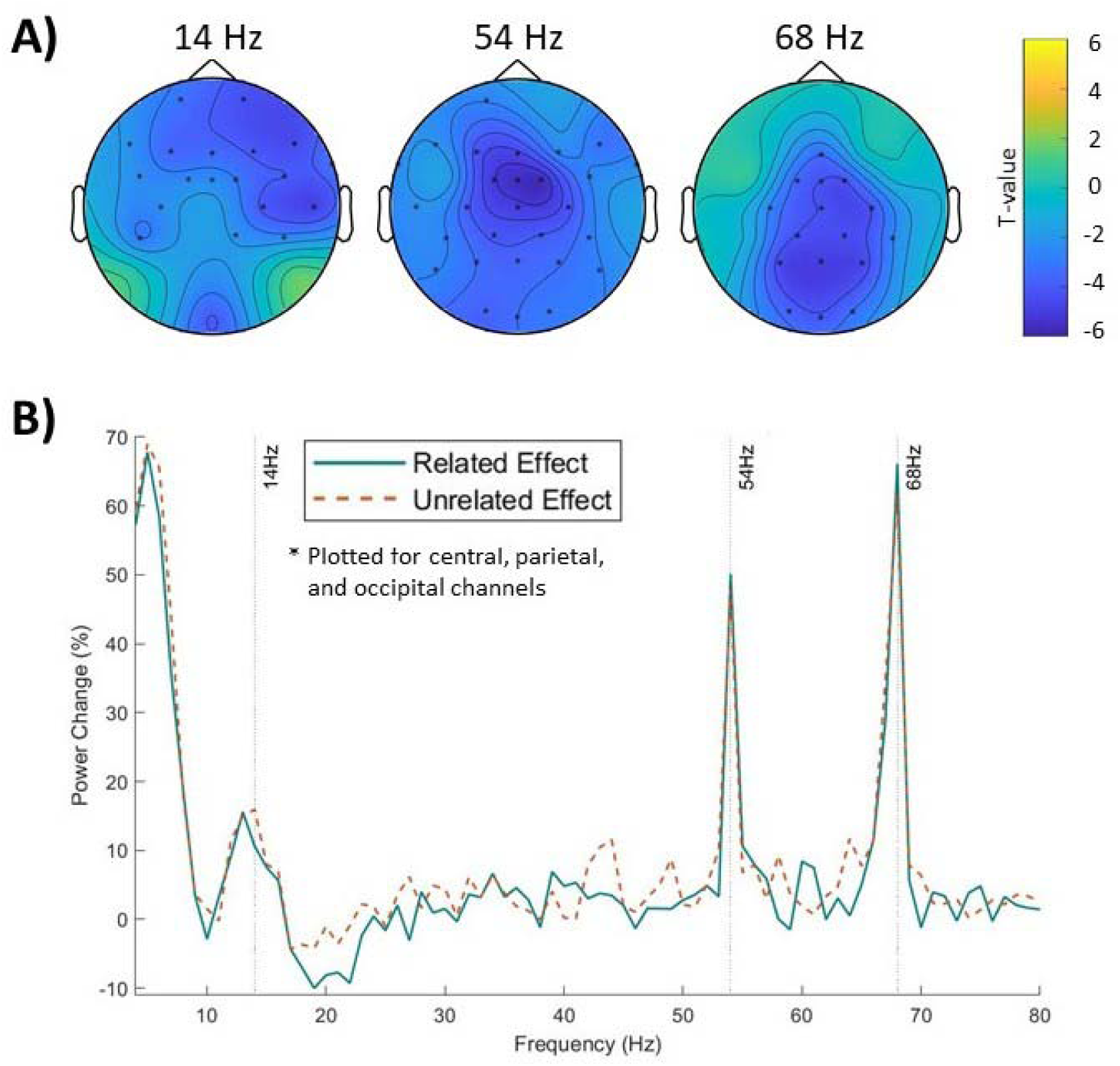
Power Analysis. A) Shows that the percentage power change between tagging window and baseline was significant at 68Hz, 54Hz, and 14Hz. The electrodes for which the contrast was significant are highlighted with asterisks. B) Shows the percentage power change plotted separately for related (green solid) and unrelated (orange dashed) condition, plotted for average of central, parietal, and occipital electrodes. The percentage power change did not differ between related and unrelated conditions at 68Hz, 54Hz, or 14Hz.

Subsequently, we computed the percentage power change between the tagging window and the baseline for both conditions (Figure 2B). The nonparametric cluster-based permutation tests found no clusters between the conditions, this shows that neither the visual SSEP or the auditory SSEP differed between the related and unrelated condition.

### 3.3. Intermodulation frequency

For the intermodulation frequency at 14Hz, we also examined whether the SSEPs were larger in the tagging window compared to the baseline. The nonparametric cluster-based permutation test showed significant clusters in frontal and central electrodes (Figure 2A) between the tagging window and the baseline (p < 0.001).

To determine which electrodes should be analyzed to examine the differences between the conditions at the intermodulation peak, we followed similar approach as Ferrante and colleagues (2023). Per participant, we selected 6 electrodes that showed the largest percentage power difference between the baseline and tagging window at 14Hz. From these electrodes, we chose the biggest cluster of electrodes based on the neighborhood structure for further analysis. For example, if all six electrodes were neighbors (based on the neighborhood structure), or in other words, electrodes in close proximity, all electrodes were included in the subsequent analysis, however, if one or two electrodes were not neighbors based on the neighborhood structure, they would be excluded from the analysis. For a sanity check, we have also followed the same electrode selection procedure at neighboring frequencies of 12Hz, 13Hz, 15Hz, and 16Hz to determine how the selection procedure in itself affects the peak.

Figure 3B shows the electrodes that were selected for the analysis at 14Hz. The majority of the channels selected for the 14Hz analysis were the left frontal electrodes, which is in line with the previous studies that found the intermodulation frequency (Drijvers et al., 2021; Seijdel et al., 2024). Subsequently, we averaged the signal over the selected electrodes based on the 14Hz effect separately for the time periods of interest (i.e., baseline and tagging window) and the conditions (related and unrelated). We then computed a dependent samples t-test between related and unrelated condition for the percentage power difference. This analysis showed two important findings. Firstly, there is a clear peak precisely at 14Hz for both related and unrelated conditions (Figure 3A). The results of the sanity check revealed that the peak at 14Hz (based on the electrode selection at 14Hz) had higher power than the peaks at the surrounding frequencies (based on the electrode selection at neighboring frequencies; for a figure see Supplementary Materials). This indicates that intermodulation frequency might manifest in different channels in EEG especially when it comes to complex paradigms that are associated with their own oscillatory pattern of results (Krott et al., 2019; Piai et al., 2012, 2014). Secondly, the t-test showed higher steady-state responses at 14Hz for the percentage power change in the unrelated compared to the related condition (t (29) = − 3.02, p = 0.005, d = 0.55), which indicates that the representations of the visual and auditory stimuli interact differently depending on their relatedness.

**Figure 3.**
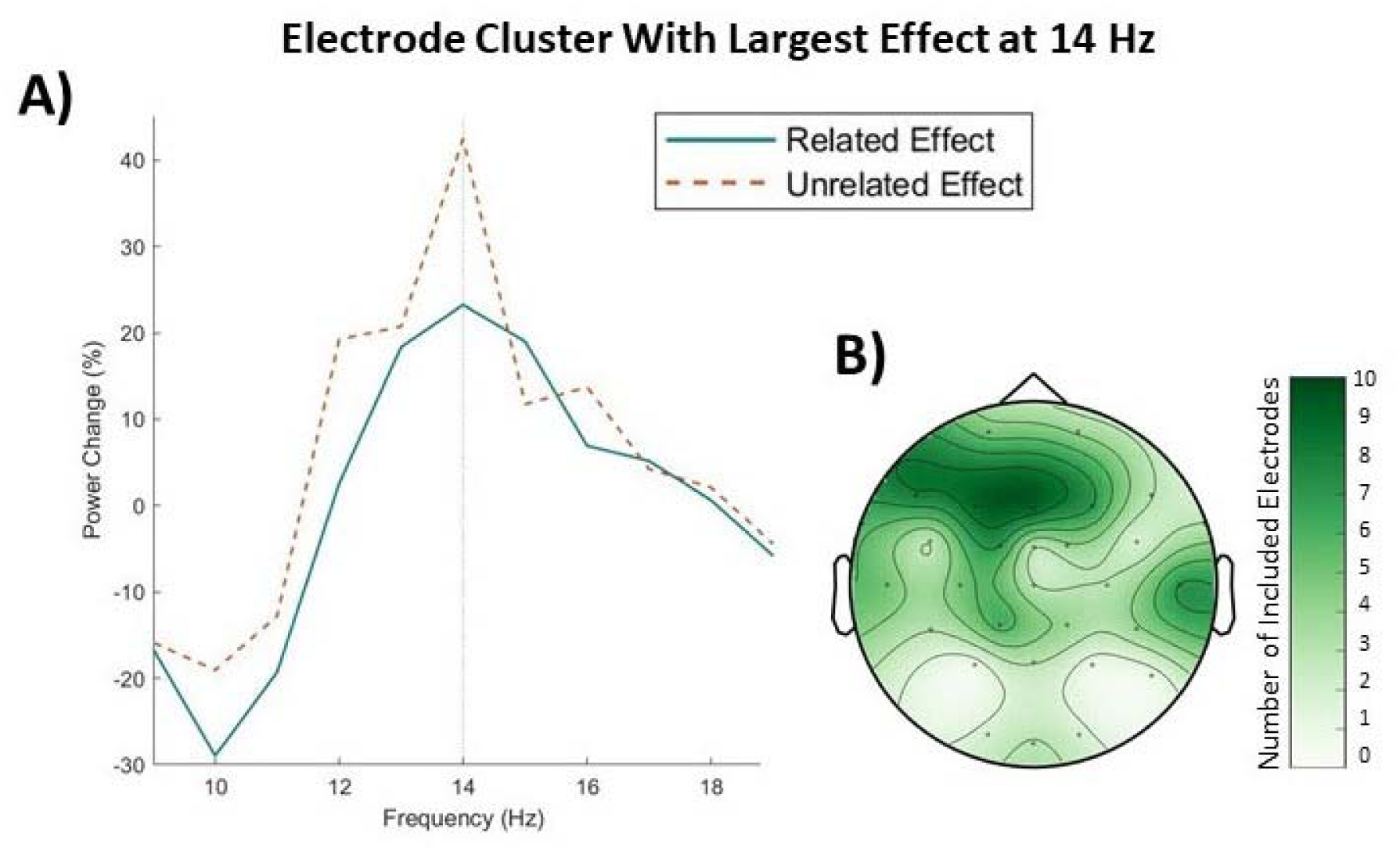
Power Analysis for Electrodes That Show Largest Effect at 14Hz. A) Shows the percentage power change (((Pow during tagging window. – Pow during baseline) / Pow during baseline) *100) plotted separately for related (green solid) and unrelated (orange dashed) condition. B) Shows the electrodes for which the power effect is plotted. The electrodes were calculated based on the largest percentage power change between tagging window and baseline (per participant) calculated for 14Hz for both conditions combined. The color scale shows the number of electrodes included at each location.

This electrode-selection approach is indeed useful, as there might be an overlap between our intermodulation frequency and the oscillatory effects elicited by the PWI paradigm (Krott et al., 2019; Piai et al., 2012, 2014). Our intermodulation results are likely not influenced by the PWI oscillatory results, especially because the previous studies found the opposite effect with respect to relatedness (higher power in related as compared to unrelated condition). If this analysis procedure was not followed, cluster-based permutation test would not reveal differences between the percentage power change effect in the related and the unrelated conditions in the steady-state responses at 14Hz (p = 0.290). Thus, this approach seems to separate the intermodulation peak from the overlapping or surrounding effects.

## 4. Discussion

In this EEG study, we utilized the RIFT method to ask (1) how people distribute their attentional resources when they are speech planning during comprehension and (2) how people’s representations of the speech planning and comprehension stimuli interact on a neural level, based on their relatedness. Our results show that the relative power change did not significantly differ between the related and unrelated condition at either 68Hz (i.e., visual tagging frequency) or at 54Hz (i.e., auditory tagging frequency). This indicates that relatedness does not affect early attention to comprehension or speech planning stimuli. Crucially, we found higher relative power change at 14Hz (i.e., intermodulation peak) in the unrelated compared to the related condition, meaning that the representations of the visual speech planning stimulus and the auditory comprehension stimulus interacted more in the unrelated condition.

### 4.1. Relatedness does not affect early attention to comprehension or speech planning stimuli

We looked at visual and auditory SSEPs to tap into the visual attention to the speech planning stimulus and the auditory attention to the comprehension stimulus, as the SSEPs reflect attention towards the tagged stimuli (Toffanin et al., 2009). We expected to find that participants would direct different amounts of attention to the visual and auditory stimuli depending on their relatedness.

Our results showed that neither the visual nor the auditory SSEPs significantly differed between the related and unrelated conditions. This indicates that the relatedness of the stimuli did not affect early visual and auditory attention. Similarly in a previous RIFT study, relatedness between words and gestures did not affect visual or auditory attention (Drijvers et al., 2021). Our results are in line with the PWI literature that posits that the interference effect arises in post-perceptual stages, either at the level of lexical selection (e.g., Levelt et al., 1999; Roelofs, 1992; Schriefers et al., 1990), which begins around 150ms after picture onset (Indefrey & Levelt, 2004), or later at the level of response selection (e.g., Finkbeiner & Caramazza, 2006; Mahon et al., 2007), which takes place after 350ms post picture onset (Indefrey & Levelt, 2004). Thus, the perceptual phase of comprehension and speech planning seems to be free from the effects of relatedness. Considering that the SSEPs reflect the perceptual attention to the comprehension and speech planning stimuli, the present results cannot speak to whether the attention to post-perceptual stages of comprehension or speech planning are affected by relatedness. In summary, our results show that auditory attention to the distractor, or early comprehension, is not affected by the relatedness of the speech planning and the comprehension content. Likewise, visual attention to the target, or early speech planning, is not affected by the relatedness.

### 4.2. Interaction between visual and auditory representations is affected by their relatedness

This study also examined the intermodulation frequency that is thought to reflect the strength of the interaction between representations of the tagged stimuli (Drijvers et al., 2021; Seijdel et al., 2024). We wanted to examine whether the interaction between the representations of the speech planning and comprehension stimuli depends on their relatedness. We wanted to test whether the interaction is stronger in the related condition, as there are more similarities between the stimuli, or rather whether the interaction is stronger in the unrelated condition, which is the easier condition, as it is associated with faster naming latencies (Bürki et al., 2020).

The central finding of this study is that the strength of the intermodulation peak depended on the relatedness of the auditory comprehension stimuli and the visual speech planning stimuli. We found a higher intermodulation peak in the unrelated compared to the related condition, meaning that the representations of the visual speech stimulus guiding speech planning and the auditory comprehension stimulus interacted more in the unrelated condition. At first glance it might seem surprising that the interaction between the representations would be stronger in the unrelated condition. However, from previous studies (as well as from our behavioral pilot) we know that naming is faster, and thus easier, in the unrelated condition (Bürki et al., 2020). This means that categorical relatedness between the stimuli makes part of the speech planning process more difficult. In order to minimize speech planning difficulties, participants might try to avoid the semantic interaction of the representations in the related condition.

It is still debated what processes give rise to and affect the intermodulation peak. Previous studies took the intermodulation frequency to represent the strength of interaction between the representations of the two stimuli (Drijvers et al., 2021; Seijdel et al., 2024). Our results indicate that if the intermodulation peak indeed represents ease of integration (i.e., stronger power reflects more integration), the intermodulation peak must also be affected by inhibitory or executive control processes that would lead to suppressed integration in the related condition. If this is the case, future studies could link the intermodulation peak to alpha power, which is thought to reflect inhibitory processing (e.g., Hilla et al., 2020; Wang et al., 2022). Correlation between the intermodulation frequency and the alpha power should be stronger in conditions that require more inhibitory processing, such as the related condition in PWI studies. An alternative explanation of the intermodulation effect could stem from the assumption that simultaneously appearing auditory and visual stimuli always interact to form a cohesive percept. The interaction could be more automatic or easier when the stimuli are related versus unrelated. Perhaps the more automatic or easier interaction could lead to smaller power in the intermodulation peak. This interpretation would mean that the intermodulation peak would reflect ease of integration, which should not be affected by inhibitory processes. If this is the case, alpha power should not be different between the related and unrelated conditions in PWI studies.

### 4.3. Implications for conversation

This study offers direct evidence that representations arising from comprehension and speech planning streams interact on a neural level. This points to the conclusion that comprehension and speech production streams interact through semantic representations. The interaction in this study was present even though processing of the stimuli in the comprehension stream was not task relevant, which speaks to the automaticity of this interaction.

The current study leaves several open questions when it comes to the interpretation of the results. For example, is speech planning crucial for the rise of the integrative process between the auditory and visual representations, or would we see the same interaction if a simple button-press task were used? The present study examined the interaction between speech planning and comprehension when both stimuli were nouns. However, in PWI studies the interference effects are affected by more factors than just semantics, such as parts of speech (Fang et al., 2015; Mahon et al., 2007). Future studies could present participants with verb distractors instead of nouns. This could shed light on whether different parts of speech affect the integrative processes as well. In PWI studies, using verb distractors tends to flip the interference effect into facilitation. Thus, using verb distractors could also help establish whether the strength of the intermodulation peak tracks the naming latencies. If this is the case, we should see higher intermodulation peak for the related than unrelated verb distractors.

### 4.4. Implications for future RIFT studies

This is the first study that combined the RIFT method with EEG (instead of MEG) in an experimental paradigm. Our results show the robustness of the SSEPs as well as the intermodulation frequency across these neuroimaging tools. In the current study the intermodulation peak was also set at a relatively higher frequency (i.e., 14Hz) than in the studies before, which used frequencies of 7Hz and below (Drijvers et al., 2021; Seijdel et al., 2024). Together, these findings show the flexibility as well as reliability of the RIFT methodology.

Importantly, this is the first study to show that semantic relatedness between the tagged stimuli alone affects the intermodulation frequency. Other studies have shown that the strength of the intermodulation peak was sensitive to the ease of lower-order audiovisual information (Drijvers et al., 2021; Seijdel et al., 2024). Our results indicate that the intermodulation peak represents more than just simple overlap between the visual and auditory representations. Instead the intermodulation peak seems to be sensitive to an interaction of the stimuli on a semantic level as well.

### 4.5. Conclusion

Our results are in line with previous behavioural work demonstrating that related distractors hinder naming pictures relative to unrelated ones (Bürki et al., 2020). Our results go beyond this work by showing that participants do not have more difficulty visually attending to the related pictures or inhibiting the related auditory distractors. Instead, processing difficulties arise when the two representations of the stimuli interact, meaning that participants might be trying to prevent integration of the auditory and visual stimuli in the related condition. This study demonstrates that RIFT is an excellent approach to investigate interactive processes between speech planning and comprehension.

