## Supplementary Materials for "Using rapid invisible frequency tagging (RIFT) to probe the attentional distribution between speech planning and comprehension"

C. Husta

Wundtlaan 1
6525 XD NIJMEGEN
NETHERLANDS

1. **Stimulus list**

|  | **Target** | **Target Translation** | **Target**  **WF** | **Related Dist.** | **Related Dist. Translation** | **Related Dist. WF** | **Unrelated**  **Dist.** |
| --- | --- | --- | --- | --- | --- | --- | --- |
| 1 | dokter | doctor | 4.0283 | verpleegster | nurse | 2.8351 | tijdschrift |
| 2 | boek | book | 3.8196 | tijdschrift | magazine | 2.6274 | verpleegster |
| 3 | bezem | broom | 2.2227 | stofzuiger | hoover | 2.0334 | aardappel |
| 4 | wortel | carrot | 2.4281 | aardappel | potato | 2.1673 | stofzuiger |
| 5 | helikopter | helicopter | 2.9814 | vliegtuig | plane | 3.5947 | computer |
| 6 | tablet | tablet | 2.0043 | computer | computer | 3.3212 | vliegtuig |
| 7 | trampoline | trampoline | 1.5315 | glijbaan | slide | 1.3617 | croissant |
| 8 | stokbrood | baguette | 1.3222 | croissant | croissant | 1.415 | glijbaan |
| 9 | olifant | elephant | 2.721 | neushoorn | rhino | 2.0414 | ingenieur |
| 10 | wetenschapper | scientist | 2.7275 | ingenieur | engineer | 2.415 | neushoorn |
| 11 | meer | lake | 4.9495 | waterval | waterfall | 2.248 | plakband |
| 12 | ducttape | duct tape | na | plakband | band aid | 2.0492 | waterval |
| 13 | stropdas | tie | 2.2945 | sjaal | scarf | 2.3674 | nieren |
| 14 | longen | lungs | 2.7324 | nieren | kidneys | 2.2577 | sjaal |
| 15 | bed | bed | 4.0209 | zitbank | sofa | 0.4771 | telraam |
| 16 | rekenmachine | calculator | 1.415 | telraam | abacus | 0.7782 | zitbank |
| 17 | paard | horse | 3.5632 | kameel | camel | 2.0755 | autobus |
| 18 | trein | train | 3.5051 | autobus | bus | 1.0414 | kameel |
| 19 | handdoek | towel | 2.6628 | toiletpapier | toilet paper | 1.9345 | dobbelsteen |
| 20 | speelkaarten | cards | 1.1461 | dobbelsteen | dice | 1.5051 | toiletpapier |
| 21 | lamp | lamp | 2.7839 | schilderij | painting | 2.9741 | portemonnee |
| 22 | rugzak | backpack | 2.5211 | portemonnee | wallet | 2.7404 | schilderij |
| 23 | frisbee | frisbee | 1.9085 | springtouw | jump rope | 0.8451 | bodywarmer |
| 24 | jas | jacket | 3.3233 | bodywarmer | bodywarmer | na | springtouw |
| 25 | oor | ear | 3.0394 | tanden | teeth | 3.2702 | prinses |
| 26 | koning | king | 3.7824 | prinses | princess | 3.269 | tanden |
| 27 | kat | cat | 3.364 | papegaai | parrot | 2.1614 | nagels |
| 28 | haar | hair | 5.1144 | nagels | nails | 2.7574 | papegaai |
| 29 | ijsblokje | ice cube | 1.2788 | sneeuwbal | snow ball | 1.6335 | schildpad |
| 30 | krab | crab | 2.2304 | schildpad | turtle | 2.2833 | sneeuwbal |
| 31 | schep | shovel | 2.3075 | kruiwagen | wheelbarrow | 1.7482 | fotograaf |
| 32 | soldaat | soldier | 3.3655 | fotograaf | photographer | 2.6375 | kruiwagen |

Table 1. Shows the stimuli used in the EEG experiment. The table includes the names of the picture targets, as well as the associated related and unrelated auditory distractors that were used. The table includes translations and word frequencies (WF) for the target and distractor names.

2. **Coherence analysis**

Coherence was calculated for every trial between the EEG signal and three dummy signals oscillating at 68 Hz, 54 Hz, and 14 Hz (Figure 1). The coherence measure was computed for frequencies between 0 and 80 Hz separately per baseline period and the tagging window using the boxcar filter and the mtmfft method. We evaluated coherence differences between the tagging window and the baseline at the main tagging frequencies (i.e. 68 Hz and 54 Hz) and at the intermodulation frequency (i.e. 14 Hz) using two-tailed non-parametric cluster-based permutation tests (Maris & Oostenveld, 2007) using 5000 permutations and the 0.05 significance threshold. The differences between related and unrelated condition were examined only after the tagging peaks significantly differed from the baseline.

Coherence at 68 Hz between the EEG signal and the 68 Hz dummy signal was significantly higher during the tagging window (p < 0.001). However, there were no clusters found when comparing the related and unrelated conditions. Similarly, coherence at 54 Hz between the EEG signal and the 54 Hz dummy signal was also significantly higher during the tagging window (p < 0.001), with no clusters found for the contrast between conditions. These results point to the same conclusions as the outcome of our main power analysis. Coherence at 14 Hz between the EEG signal and the 14 Hz sine wave was not significantly different between the baseline and the tagging window (p = 0.09). This is not surprising, as previous studies showed that the intermodulation peak does not manifest in coherence, but rather in power (Drijvers et al., 2021) especially when the moment of interaction might differ.

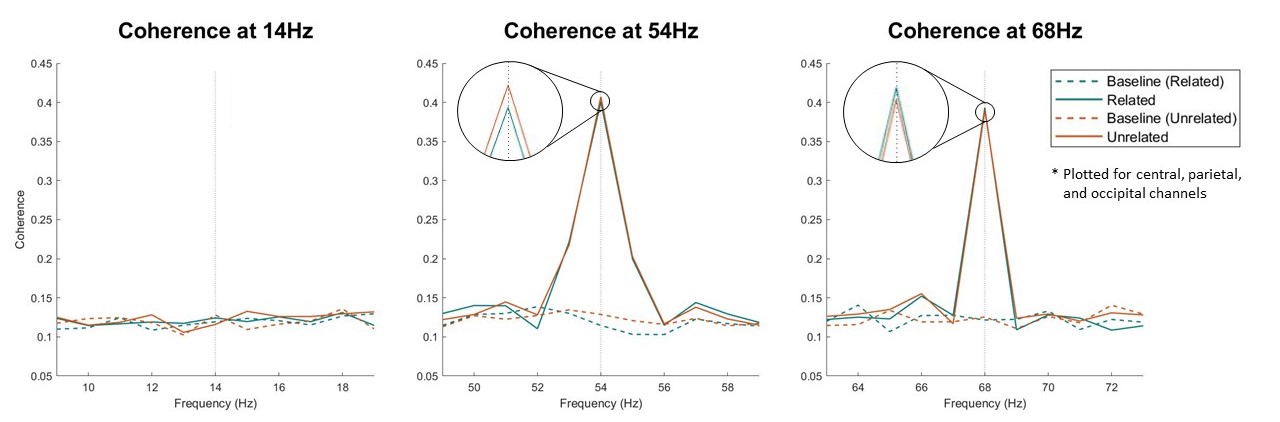

***Figure 1. Coherence Analysis.*** *The plots show coherence between the EEG signal and 14 Hz (left plot), 54 Hz (middle plot), and 68 Hz (right plot) for the central, parietal and occipital channels. The plots include coherence during the baseline period (dashed lines) as well as during the tagging window (solid lines) for the related (green) as well as unrelated condition (orange).*

3. **Validation of the electrode selection procedure**
   3.1. Selection of electrodes for the main tagging frequencies

As this is one of the first studies using RIFT with EEG, the standard analysis protocol has not yet been established. However, previous studies have analyzed RIFT data for different sensors per participant, depending on which sensor showed the strongest tagging response (Ferrante et al., 2023). Following Ferrante et al., 2023, we used a similar procedure to identify the intermodulation peak. Per participant, we selected 6 electrodes that showed the largest percentage power difference between the baseline and tagging window. From these electrodes, we chose the biggest cluster of electrodes based on the neighborhood structure for further analysis. For example, if all six electrodes were neighbors, or in other words electrodes in close proximity, all electrodes were included in the subsequent analysis, however, if one or two electrodes were located far away from the others, they would be excluded from the analysis. To establish whether this electrode selection procedure works, we have first performed it for the effects at 54 Hz and 68 Hz. Figure 2 shows that this analysis procedure selects mainly central electrodes for the 54 Hz effect and mainly occipital electrodes for the visual 68 Hz effect. Effects in these locations correspond well to where the auditory and visual effects should manifest in EEG. Thus, this electrode selection procedure could be used to capture the electrode cluster where the intermodulation peak manifests per participant.

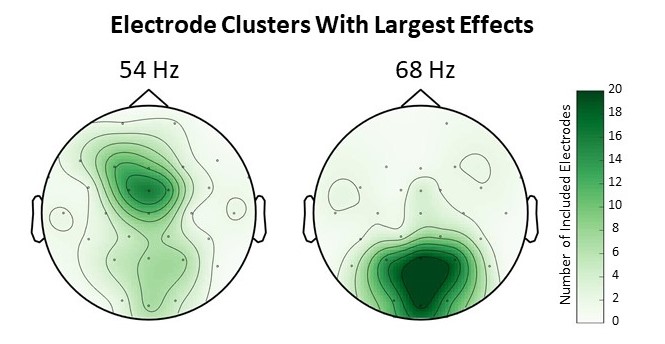

***Figure 2. Electrodes That Show Largest Effect.*** *This plot shows the electrodes for which the power effect (((Pow during tagging window. – Pow during baseline) / Pow during baseline) *100) was the largest per participant at 54 Hz (left plot) and at 68 Hz (right plot). The color scale shows the number of electrodes included.*

3.2. Selection of electrodes at frequencies neighboring 14Hz

In the main manuscript we describe that the electrode selection procedure at 14 Hz results in a better-defined intermodulation peak. However, we wanted to establish how stable the intermodulation effect is, if the electrodes would be selected based on the biggest power change in neighboring frequencies. Thus, per participant, we computed electrode clusters that showed biggest power change at 12 Hz, 13 Hz, 15 Hz, and 16 Hz. We then plotted power change for each of these electrode selections. Figure 3 shows that the highest peak results at 14 Hz for the electrode cluster selected based on the largest difference between baseline and tagging window at 14 Hz. This supplementary analysis shows that employing a procedure to find electrodes where the intermodulation peak is most pronounced might be necessary when the intermodulation peak is investigated with EEG or in a paradigm associated with its own pattern of potentially overlapping oscillatory results.

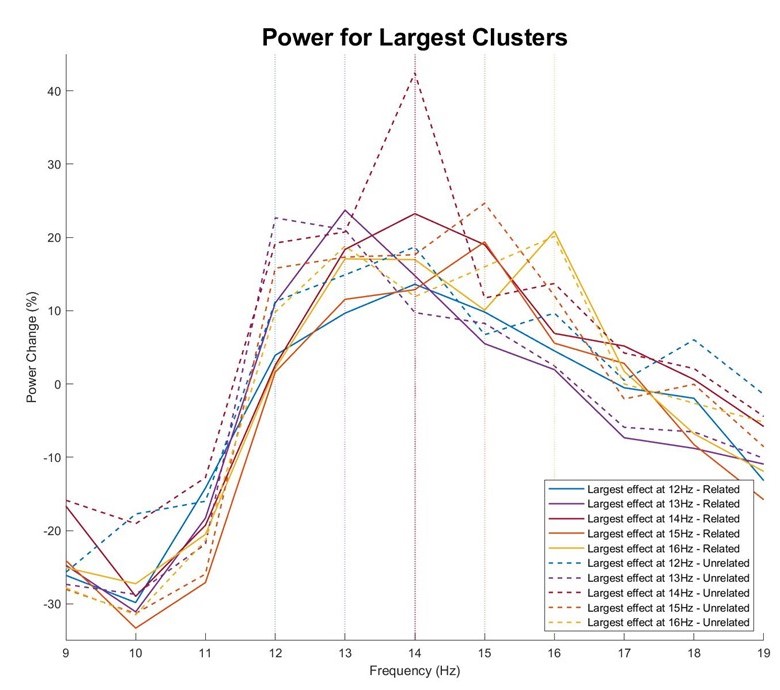

***Figure 3. Power Analysis for Electrodes That Show Largest Effect at 12 Hz, 13 Hz, 14 Hz, 15 Hz, and 16 Hz.*** *This plot shows the percentage power change (((Pow during tagging window. – Pow during baseline) / Pow during baseline) *100) plotted separately for related (solid) and unrelated (dashed) condition. The percentage power change is plotted for clusters of electrodes that show largest effects between baseline and the tagging window at 12 Hz (blue), at 13 Hz (purple), at 14 Hz (red), at 15 Hz (orange) and at 16 Hz (yellow).*
